# Cryo-EM structure of *Escherichia coli* σ^70^ RNAP and promoter DNA complex revealed a role of σ non-conserved region during the open complex formation

**DOI:** 10.1101/256826

**Authors:** Anoop Narayanan, Frank S. Vago, Kungpeng Li, M. Zuhaib Qayyum, Dinesh Yernool, Wen Jiang, Katsuhiko S. Murakami

## Abstract

First step of gene expression is transcribing the genetic information stored in DNA to RNA by the transcription machinery including RNA polymerase (RNAP). In *Escherichia coli*, a primary σ^70^ factor form the RNAP holoenzyme to express housekeeping genes. The σ^70^ contains a large insertion at between the conserved regions 1.2 and 2.1, the σ non-conserved region (σ_NCR_), but its function remains to be elucidated. In this study, we determined the cryo-EM structures of the *E. coli* RNAP σ^70^ holoenzyme and its complex with promoter DNA (open complex, RPo) at 4.2 and 5.75 Å resolutions, respectively, to reveal native conformations of RNAP and DNA. The RPo structure presented here found an interaction between R157 residue in the σ_NCR_ and promoter DNA just upstream of the −10 element, which was not observed in a previously determined *E. coli* RNAP transcription initiation complex (RPo plus short RNA) structure by X-ray crystallography due to restraint of crystal packing effect. Disruption of the σ_NCR_ and DNA interaction by the amino acid substitution (R157E) influences the DNA opening around the transcription start site and therefore decreases the transcription activity of RNAP. We propose that the σ_NCR_ and DNA interaction is conserved in proteobacteria and RNAP in other bacteria replace its role with a transcription factor.

## INTRODUCTION

In the event of formation of the transcription ready open complex (RPo) in *Escherichia coli*, the conserved regions 2.4 and 4.2 of the primary σ^70^ factor in RNA polymerase (RNAP) holoenzyme recognize the −10 and −35 promoter elements, respectively, to form the closed complex (RPc). This step is followed by DNA strands separation that is stabilized by the σ region 2.3, completing the formation of the transcription bubble from −11 to +2 positions (1). *E. coli* RNAP forms RPo with at least two intermediates starting from the RPc. Although a series of large conformation changes of RNAP and DNA during the RPo formation have been characterized by kinetic and biophysical studies (2), the structures of intermediates have not been determined yet due to their transient and heterogeneous nature that make them difficult to be captured by X-ray crystallography approach.

A series of technical advances such as development of a direct electron detector and also image processing techniques have moved cryo-electron microscopy (cryo-EM) to the forefront of structural biology to analyze large macromolecule complexes in unprecedented details (3). Several structures of *E. coli* RNAP in complex with nucleic acids (DNA/RNA and 6S RNA) (4,5) and transcription activators (NtrC and CAP) (6,7) have been determined by cryo-EM, demonstrating the feasibility of using this experimental approach to investigate the structure and function of *E. coli* RNAP. There are a couple of advantages of the cryo-EM over X-ray crystallography: 1) the 3D classification of individual particles from a single cryo-EM grid makes it possible to reveal the heterogeneity of macromolecule and determine their structures individually; 2) a cryo-EM grid preparation takes less than 10 sec instead of taking days or longer for macromolecular crystallization, increasing our chances to capture elusive and unstable intermediates for cryo-EM structure determination.

The structural study of bacterial RNAP began in 1996 with the high-resolution X-ray crystal structure of the *E. coli* σ^70^, which provided insight into the DNA sequence recognition and double-strand DNA melting by the σ factor conserved regions 2.4 and 2.3, respectively (8). The structure also revealed the s non-conserved region (σ_NCR_) at between the conserved regions 1.2 and 2.1.

In this work, we determined the cryo-EM structures of the *E. coli* RNAP σ^70^ holoenzyme and the holoenzyme-promoter DNA complex from a single cryo-EM grid. Comparing these structures revealed that the promoter DNA binding to RNAP not only triggers conformational changes of RNAP around the domains for binding the −35 and −10 elements, but also establishes the direct interaction between the σ_NCR_ and the DNA upstream of the −10 element. Our structural and biochemical data introduced a function of the σ_NCR_ during the formation of transcription ready RPo.

## RESULTS

### Sample preparation and cryo-EM structure determination

In this study, we aimed to determine the structure of *E. coli* RNAP σ^70^ holoenzyme and promoter DNA complex with KdpE (KDP operon transcriptional regulatory protein) to obtain structural basis of the transcription activation by the response regulator OmpR/PhoB family. We assembled a ternary complex (**Fig. S1**) containing *E. coli* RNAP, a constitutively active variant of KdpE (KdpE-E216A, (9)) and the *E. coli kdpFABC* promoter DNA derivative containing both a RNAP binding site (−35 and −10 elements; synthetic bubble from −6 to +1) and tandem KdpE binding sites (from −67 to −62 and from −56 to −51) (pFABC-1, **Fig. S2**) for the cryo-EM grid preparation.

Out of many grid types on which the sample was vitrified, lacey-carbon grids coated with a single layer of graphene oxide followed by a layer of pyrene-NTA had images with more widely oriented particles, better suited for single particle 3-D reconstruction. From 1,037 usable movie stacks, 286,424 particles were picked and used for single particle reconstruction. The 54 classes with clear structural features representing 257,539 particles after 2D classification were selected for 3D classification. This resulted in two major 3D classes, one representing an *E. coli* RNAP σ^70^ holoenzyme-promoter DNA complex (RPo) and the other of *E. coli* RNAP σ^70^ holoenzyme by itself (**Fig. S3**) indicating that the dissociation of DNA from some fractions of the RPo during the cryo-EM grid preparation. We failed to obtain a density map representing a ternary complex (RPo with KdpE), suggesting the dissociation of KdpE from the RPo. Determination of the cryo-EM structures of the RPo and the apo-form RNAP from a single cryo-EM grid allows us to reveal conformational changes of RNAP upon the promoter DNA binding in the identical experimental condition in solution.

### Cryo-EM structure of the E. coli RNAP open complex

One of the 3D classes represents the *E. coli* RPo determined at 5.75 Å resolution (**Figs. 1** **and S3**). In addition to density for RNAP holoenzyme, strong densities are observed for double-stranded DNA (from −45 to −12 bases) and single-stranded non-template DNA (nt-DNA) in the transcription bubble (from −11 to −3 bases). However, density for the template DNA (t-DNA) in the transcription bubble (from −9 to +2 bases) is not traceable and density for downstream DNA duplex is weak and scattered, indicating their mobility within the DNA binding main channel of RNAP (**Fig. 1C**). Positions of each DNA elements (−35, −10, transcription bubble and downstream DNA) are nearly identical to a previously determined X-ray crystal structure of *E. coli* transcription initiation complex (TIC) (10) (PDB: 4YLN, **Fig. S4A**). Although the overall resolution was 5.7 Å, local resolution calculations indicate the central part of the structure including the N-terminal domain of α subunit (αNTD) as well as β and β’ subunits around the active site of RNAP was determined close to 5 Å resolution, with peripheral areas of the structure such as σ_NCR_ (N128 to R373, 246 residues), β’ insertion 6 (β’i6, A944 to G1129, 186 residues) and ω subunit at 7-10 Å resolution (**Fig. 1D**). Density for the C-terminal domains of α subunit (αCTD) and σ region 1.1 domain (σ_1.1_) could not be traced due to flexible nature of these domains. The distance between two pincers of RNAP (T212 in the β’ clamp domain and H165 in the β gate loop) is 28 Å, indicating the closed conformation of the RNAP clamp as observed in the *E. coli* TIC (27.7 Å). However, the position of σ_NCR_ in the RPo structure determined by the cryo-EM in this study is different from the one in the TIC determined by the X-ray crystallography (**Fig. S4A**). Particularly, the σ_NCR_ contacts the t-DNA at −16/17 position via R157 residue, while this interaction was not observed in the TIC due to restraint of crystal packing effect (**Fig. S4B**).

**Figure 1.**
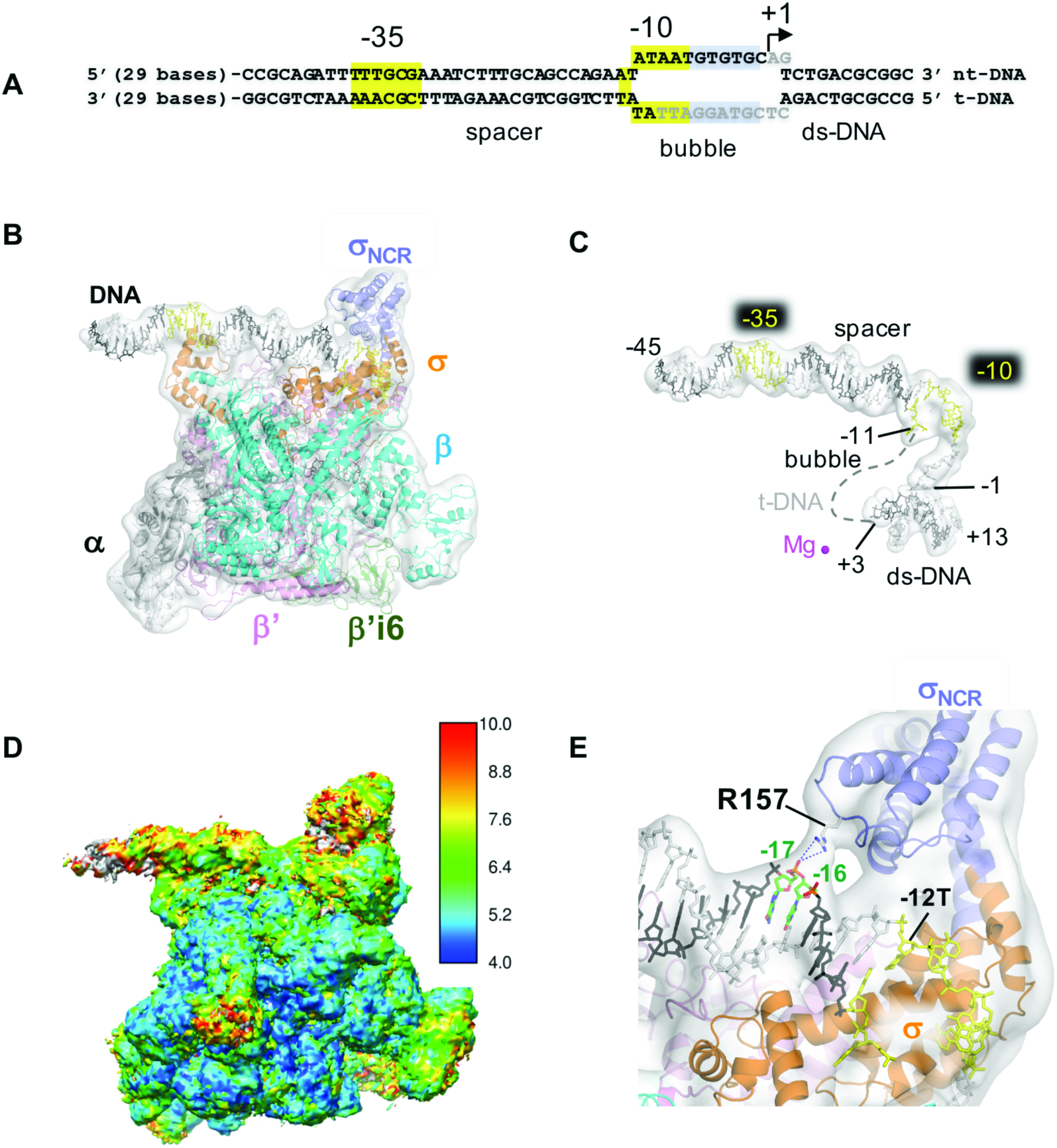
The cryo-EM structure of the E. coli RPo. *A*, Sequence of the promoter DNA visible in the cryo-EM structure of RPo (from −45 to +13) is depicted (full length DNA sequence used for the RPo preparation is shown in **Fig. S2**). −35 and −10 elements (highlighted in yellow), transcription bubble, and transcription start site (+1) are indicated. Bases shown in gray are disordered in the RPo structures. ***B***, Cryo-EM structure of RPo. Ribbon model of RNAP and stick model of DNA superimposed with the cryo-EM density map (white and transparent). Each subunit of RNAP, the σNCR (light blue) and P’i6 (green) are indicated in color. Template (t-DNA) and non-template DNA strands (nt-DNA) are depicted as dark gray and light gray, respectively and the −35 and −10 elements are shown in yellow. ***C***, Promoter DNA in the RPo. Stick model of promoter DNA with its cryo-EM density map are shown as the same orientation as in ***B***. Dashed line represents the disordered t-DNA in the transcription bubble. The active site Mg^2+^ ion is shown as magenta sphere. Nucleotide bases at −11 and +3 (representing boundaries of the transcription bubble) and transcription start site (+1) are indicated. ***D***, Cryo-EM map of RPo colored by local resolution oriented as the same orientation as in ***B. E***, A zoomed-in view of ***B*** shows the interaction between R157 residue and t-DNA at −16/−17 positions.

### Cryo-EM structure of the E. coli RNAP σ^70^ holoenzyme

The second 3D class represents the apo-form *E. coli* RNAP σ^70^ holoenzyme with an overall resolution of 4.2 Å (**Figs 2A** **and S3**). Local resolution calculations indicate that the center part of the structure is determined at a resolution of 4 Å, while the peripheral areas of the structure are determined around 6-10 Å (**Fig 2B**). Densities for αCTD and σ_1.1_ were not traceable. Densities for σ_4_, σ_NCR_ and β flap tip helix are sparse and weaker than their counterparts in the RPo, indicating their flexible nature before binding to promoter DNA. The distance between the two pincers around the main channel is 26 Å, indicating that the RNAP clamp adopts a closed conformation without DNA binding.

Density around the active site located at the center of the RNAP molecule showed not only main chains but also side chains. For example, the density map of the Rifampin binding pocket of β subunit shows main chain as well as the side chains of H526 and S531 residues that play key roles in the RNAP and Rifampin interaction (11) (**Fig. 2C**). The cryo-EM map shown here is the same quality as the X-ray crystal structure of *E. coli* RNAP determined at 3.6 Å resolution (PDB: 4YG2) (12), indicating that cryo-EM is able to provide the structural information for the RNAP and inhibitor interaction.

**Figure 2.**
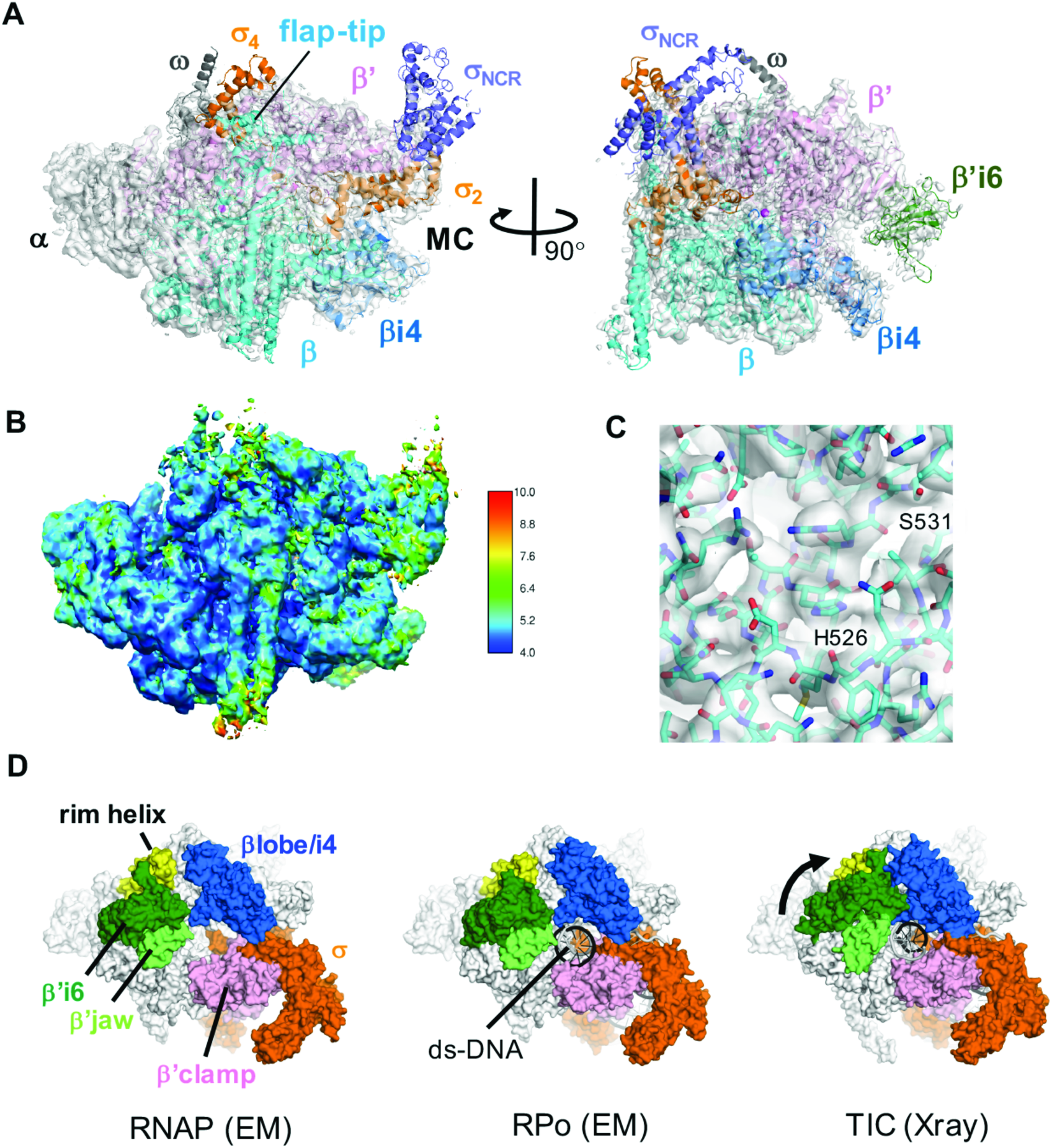
The cryo-EM structure of the *E. coli* RNAP σ^70^ holoenzyme. ***A***, Cryo-EM structure of RNAP is shown with the cryo-EM density map (white and transparent). subunits and domains of RNAP are shown as in **Fig. 1B**. A position of the RNAP main channel (MC) is indicated. ***B***, Cryo-EM map of RNAP colored by local resolution. ***C***, Rifampin binding pocket of the β subunit is shown with the cryo-EM map. S531 and H526 residues are labelled. ***D***, Comparison of the cryo-EM structures of the holoenzyme (left, this study), RPo (middle, this study) and the crystal structure of TIC (right, PDB: 4YLN). Structures are shown as surface models with σ^70^ (orange), βlobe/i4 (blue), rim helix (yellow), β’i6 (dark green), β jaw (pale green) and P’clamp (magenta). Closure of the gap between the P’i6 domain and the plobe/i4 domain observed in the TIC structure compared to others is shown by a black arrow.

In the cryo-EM structures of both apo-form RNAP and RPo, a gap between the β’i6 domain and β’ rim helix is widely opened. In comparison, the β’i6 shifts toward the β’ rim helix about 31 Å in the X-ray crystal structure of TIC, showing the complete closure of the downstream DNA binding cleft of RNAP (**Fig. 2D**). Significance of the movement of β’i6 will be discussed below.

### Cryo-EM structure reveals a novel interaction between σ_NCR_ and DNA upstream of the −10 element

Although densities of the σ_4_ and σ_NCR_ domains are blurred in the apo-form RNAP, the RPo showed well-ordered these domains. Interestingly, the σ_NCR_ is involved in the binding of upstream DNA of the −10 element. R157 residue of the σ_NCR_ establishes a long-range electrostatic interaction with a phosphate between the −16/−17 DNA positions (**Fig. 1E**).

To investigate a role of the interaction between the σ_NCR_ and promoter DNA, we prepared an *E. coli* RNAP containing the σ^70^-R157E substitution and tested its transcription activity with a double-stranded promoter DNA (pFABC-3, **Fig. S2**). The *E. coli* RNAP derivative showed major defect in transcription, expressing only 22 % activity compared with the wild-type enzyme (**Figs. 3A and B**). To identify the step of transcription influenced by the R157E substitution, we investigated the promoter DNA binding of RNAP by using Electrophoretic Mobility Shift Assay (EMSA) and found no effect by the substitution (**Fig. 3C**). To test the effect of the R157E substitution in DNA unwinding, we used fluorescence signal of 2-aminopurine (2-AP) substitutions at −7 and −1 positions in double-stranded promoter DNA (pFABC_2AP-1 and pFABC_2AP-7, **Fig. S2**).

**Figure 3.**
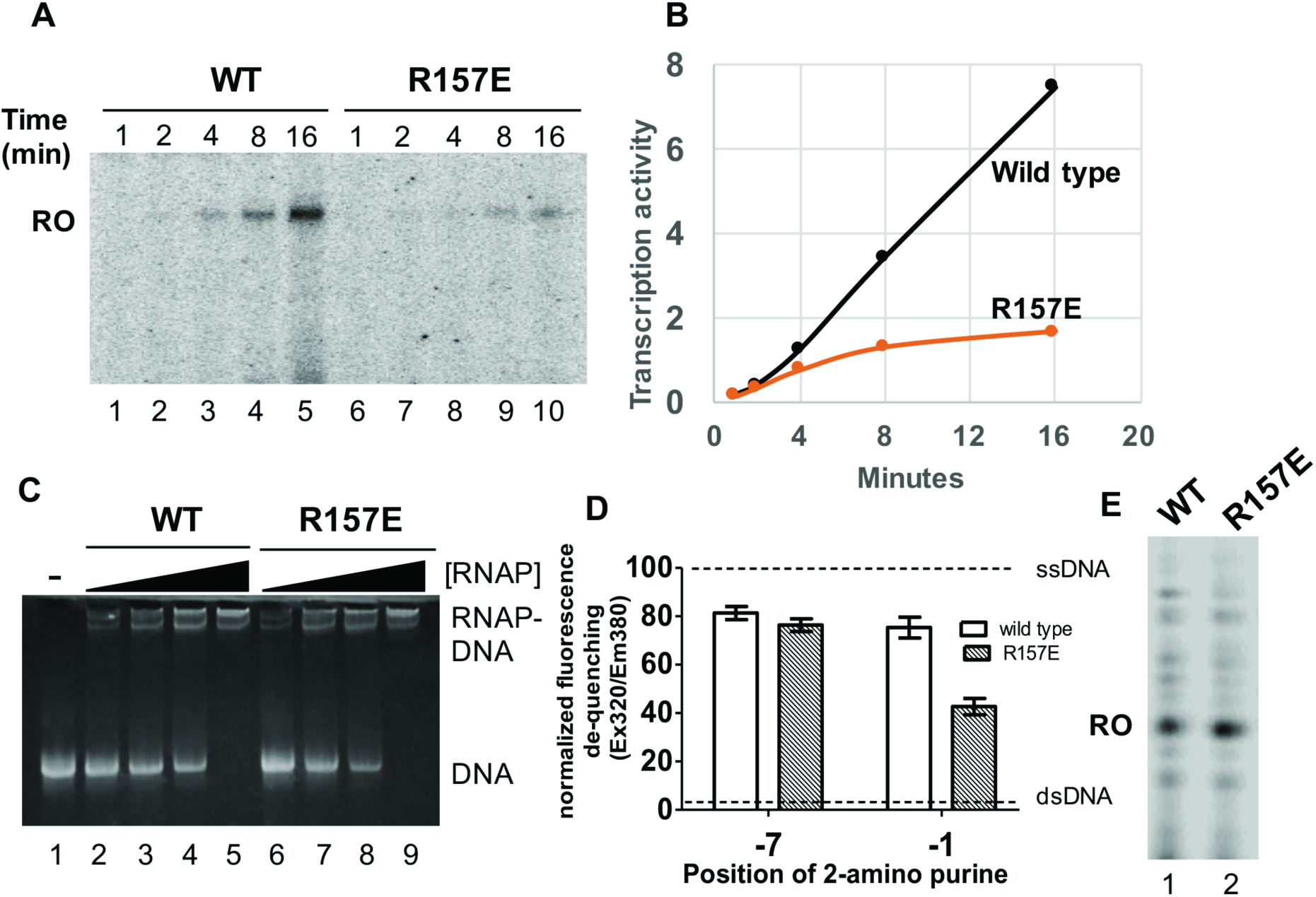
Characterization of the σ _NCR_ and promoter DNA interaction for RNAP transcription activity. ***A***, R157E substitution of σ^70^ decreases the transcription activity of RNAP. Transcription activities of wild type RNAP (lanes 1-5) and σ^70^-R157E substituted RNAP (lanes 6-10) on a double stranded promoter DNA (pFABC-3, SFig. 2) at different time intervals as indicated. Position of the run-off transcription product (RO) is shown. ***B***, Plot comparing the rate of transcription activity between the wild type and the σ^70^-R157E substituted RNAPs. The run-off transcript levels from ***A*** were quantified using the software ImageJ. ***C***, Electrophoretic Mobility Shift Assay (EMSA) showing binding of wild type (WT, lanes 2-5) and the σ^70^-R157E substituted RNAPs (R157E, lanes 6-9) to pFABC-3 promoter DNA. Lane 1 is free DNA. Positions of free DNA and RNAP-DNA complex are indicated. ***D***, Fluorescence signal from the 2-aminopurines substituted at −1 and −7 positions of the template strand promoter DNA (pFABC_2AP-1 and pFABC_2AP-7, **Fig. S2**) by wild type RNAP and σ^70^-R157E substituted RNAPs. Fluorescence data was normalized considering the difference in the fluorescence values of single stranded DNA (SsDNA, dotted line at the top) and double stranded DNA (DsDNA, dotted line at the bottom) as 100 %. Average values from three experiments is plotted using the software GraphPad Prizm. ***E***, Transcription activities of wild type and σ^70^-R157E substituted RNAPs on a pre-melted DNA scaffold (TIC scaffold, **Fig. S2**).

Fluorescence signal of 2-AP is quenched in double-stranded DNA, whereas the increase of 2-AP fluorescence intensity is observed when RNAP unwinds double-stranded DNA (13). Both wild type and derivative RNAP show similar increase of the 2-AP fluorescence intensity at −7 position, suggesting that the R157E substitution does not influence the early step of DNA opening. However, 2-AP fluorescence at −1 position was reduced by 2-folds in the derivative compared with the wild-type (**Fig. 3D**), indicating that the σ_NCR_ plays a role in the DNA unwinding around the transcription start site, the final step of transcription ready RPo formation. Consistent with this functional role, the R157E substitution had no effect (**Fig. 4E**) on the transcription of a pre-melted DNA (TIS, **Fig. S2**).

**Figure 4.**
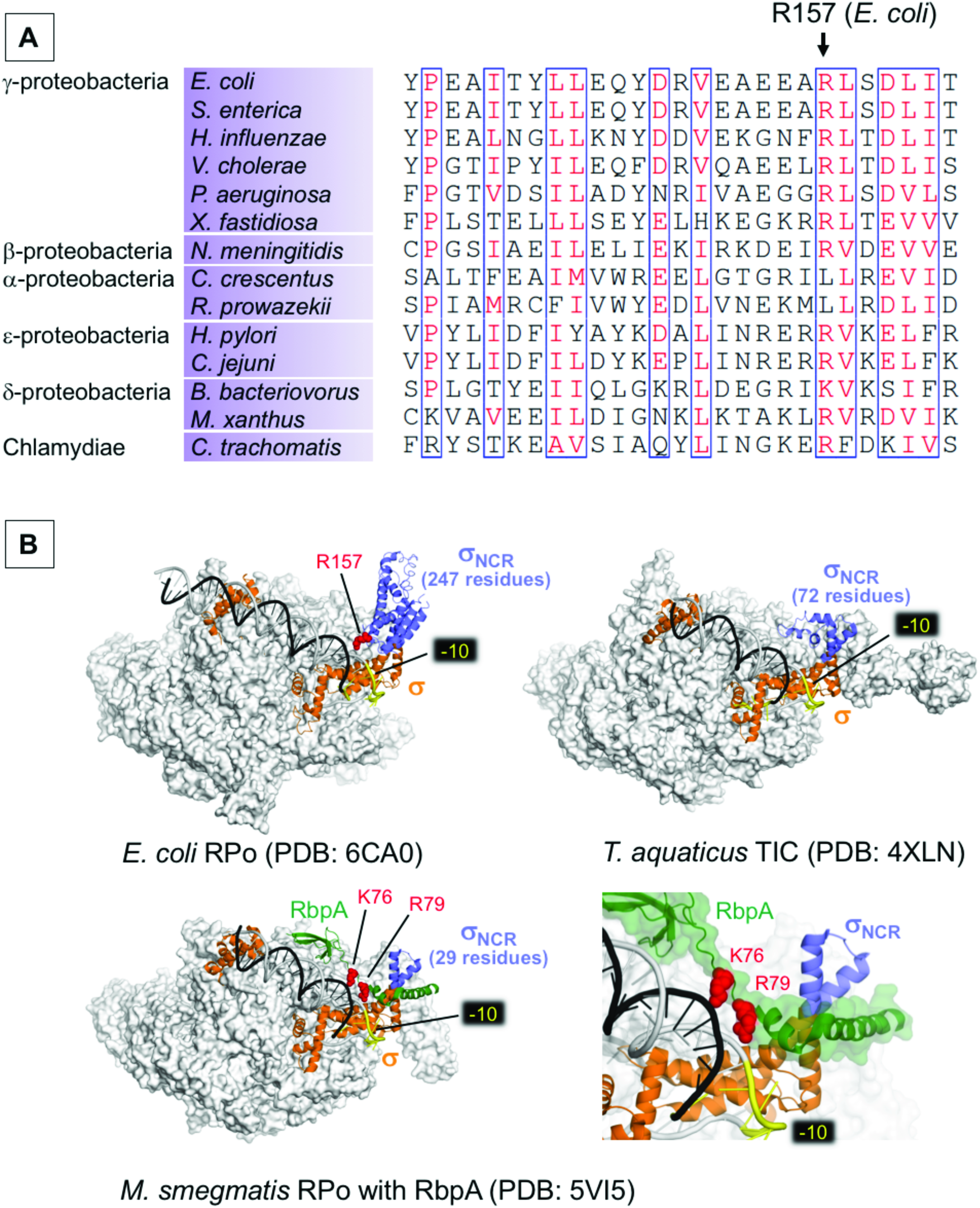
***A***, Alignment of the sigma_NCR_ region shows the conservation of R157 residue in bacteria from the classes of proteobacteria and Chlamydiae. ***B***, the structures of *E. coli* RPo (top left), *Thermus aquaticus* RPo (top right) and *Mycobacterium smegmatis* (RPo with RbpA, bottom) as transparent surface models with corresponding σ factors (orange) and promoter DNA (t-DNA-dark gray, nt-DNA-light gray) shown as ribbon models. σ_NCR_ is in purple and RbpA of *M. smegmatis* RPo (bottom) is in green. −10 elements are colored yellow. Positively charged residues interacting with bases upstream to −10 promoter elements (R157 of *E. coli σ*_NCR_, R79 and K76 of RbpA) are shown as sphere model (red). A right bottom panel is a magnified view showing the RbpA and DNA interaction.

## DISCUSSION

Using single-particle cryo-EM reconstruction, we determined the structures of the apo-form *E. coli* RNAP σ^70^ holoenzyme and RPo at 4.2 Å and 5.7 Å resolution, respectively. Although the overall resolution of the apo-form RNAP is better than RPo, the density maps of RNAP involved in the −35 element recognition such as σ_4_ and β flap tip helix are better resolved in the RPo (**Fig. 1B**, **2A**), suggesting that RNAP needs its flexibility of the −35 element binding domain to recognize promoters with different lengths of spacer (16 to 18 bases in most case) between the −10 and −35 elements.

The density map of σ_NCR_ in the apo-form holoenzyme is weak and sparse, whereas the σ_2_, which directly links to the σ_NCR_ via two a helixes, is well resolved (**Fig. 2A**). The promoter DNA binding to RNAP enhances the rigidity of σ_NCR_ (**Fig. 1B**) and establishes a long-range electrostatic interaction between R157 residue and t-DNA strand at −16/−17 position (**Fig. 1E**). This interaction was not observed in a previously determined *E. coli* RNAP TIC determined by the X-ray crystallography (10) likely due to restraint of protein and DNA packings in the crystal (**Fig. S4B**). The σ_NCR_ and t-DNA interaction presented here is distinct from the σ_3_ and DNA interaction, which is formed with the nt-DNA strand upstream of the −10 element (14).

Our structure-based biochemical experiments revealed a role of the σ_NCR_ and t-DNA interaction, which facilitates the DNA opening of the transcription start site and formation of the transcription-ready RPo (**Fig. 3**). R157 residue is widely conserved in Betaproteobacteria, Gammaproteobacteria, Epsilonproteobacteria, Deltaproteobacteria and *Chlamydiae* (**Fig. 4A**), further supporting the importance of the σ_NCR_ and DNA interaction.

The size of σ_NCR_ found in the primary σ factor (σ^70^ in *E. coli* and SigA in other bacteria) depends on bacterial phyla. For examples, *E. coli σ*^70^ contains 247 residues, whereas *Thermus aquaticus*, *Mycobacterium smegmatis* and *Mycobacterium tuberculosis* (MTB) SigA contain 72, 29 and 32 residues, respectively (**Fig. 4B**). There is a good correlation between the size of σ_NCR_ and the stability of RPo. Compared to MTB RNAP that forms very unstable RPo with two promoters tested, *E. coli* RNAP formed highly stable and irreversible complexes with the same promoters (15). Stable RPo formation of MTB RNAP requires accessary proteins RbpA and CarD (16). Due to its small size, the σ_NCR_ of MTB and *M. smegmatis* SigA cannot touch DNA in the RPo (16,17). However, in the presence of RbpA, the basic linker (BL) and σ interacting domain (SID) of RbpA occupies a space between the promoter DNA and σ_NCR_, acting like an extension of σ_NCR_ to reach DNA upstream of the −10 element (16). Basic residues (K76 and R79) of the RbpA-BL form salt bridges with phosphates of the nt-DNA upstream of the −10 elements (**Fig. 4B**) and particularly, R79 and DNA interaction is critical for the stable RPo formation (16). Thus, it is likely that the RpbA in MTB is a functional counterpart of the σ_NCR_ in the *E. coli* RNAP transcription system.

The β’i6 is a lineage specific insertion found between the middle of the trigger loop and changes its position at deferent stages in the transcription cycle. The β’i6 domain is in the open-conformation in the cryo-EM structures of RNAP presented here whereas it is in the closed-conformation in the X-ray crystal structure of TIC, respectively (**Fig. 2D**). The movement of the β’i6 domain toward the βlobe/i4 domain results in closure of the gap between the β’jaw and βlobe/i4 domains. The β’jaw and β’i6 form the downstream mobile element (DME) of RNAP and the kinetic studies of *E. coli* RNAP transcription proposed the conformational change of DME during the stable RPo formation (2,18). Deletion of β’i6 drastically reduced the stability of the RPo (19), which is consistent with the idea that the β’i6 may tighten the grip of RNAP on the downstream DNA at the late stage of RPo formation. Although both the cryo-EM and X-ray crystallographic studies used the same sequence and length of downstream DNA to prepare the RNAP-DNA complex, the downstream DNA in crystal forms longer DNA as a result of head-to-tail binding with the upstream DNA of adjacent symmetrically related molecule (**Fig. S4C**). It is tempting to speculate that certain length of downstream DNA accommodated in the DNA binding main channel triggers the conformational change of the DME for the establishment of the stable RPo formation.

## CONCLUSION

The cryo-EM structure of RNAP holoenzyme was determined at 4.2 Å resolution and its density map quality is equivalent to the X-ray crystal structure of *E. coli* RNAP holoenzyme determined at 3.6 Å resolution, indicating that single-particle cryo-EM is an alternative and promising method for structural studies of RNAP inhibitors due to eliminating the crystallization step. The cryo-EM structure of RPo presented here revealed a novel interaction between σ_NCR_ and DNA upstream of the −10 element, which facilitates the formation of stable RPo. RNAP derivative containing σ^70^-R157E accumulates intermediate species between the closed and open complexes, therefore, it, along with the cryo-EM, could be a useful tool to capture elusive intermediates during the RPo formation. Such experiments are underway.

## EXPERIMENTAL PROCEDURES

### Protein expression and purification

*E. coli* RNAP core enzyme and σ^70^ proteins were prepared and RNAP holoenzyme was reconstituted as described (20). R157E mutation in the *E. coli rpoD* gene was obtained by site directed mutagenesis of plasmid pGEMD, and σ^70^ derivative was prepared and holoenzyme containing σ^70^ derivative was reconstituted as described (20). Response regulator KdpE-E216A was expressed and purified as described (9).

### Sample preparation for cryo-EM

*E. coli* RNAP σ^70^ holoenzyme, response regulator KdpE-E216A and synthetic DNA (pFABC-1, **Fig. S2**) were mixed at 1:2:3 molar ratio in sample buffer (10 mM Hepes pH 8, 100 mM NaCl, 5% glycerol, 10 mM MgCl_2_) and incubated at room temperature for 30 minutes. The ternary complex was purified using size exclusion superose 6 column chromatography (**Fig. S1**) equilibrated with buffer containing 10 mM Hepes pH 8, 100 mM NaCl, 5% glycerol, 1 mM MgCl_2_. The ternary complex in peak fractions were pooled and cross-linked with 0.1 mM glutaraldehyde for 30 min at room temperature. The cross-linked sample showed that the RNAP subunits formed a single band representing a large crosslinked complex, while the mobility of KdpE-E216A remains the same as the non-cross-linked sample (**Fig. S1D**) suggesting that KdpE-E216A does not directly interact with RNAP. After crosslinking, buffer was exchanged to reduce glycerol concentration to < 1% and the sample was applied to lacey-carbon grids coated with a layer of graphene oxide followed by a layer of pyrene NTA.

### Grid preparation for cryo-EM

Graphene oxide solution (10 μg/ml, 3 μl) was applied to a lacey carbon grids (Ted Pella, Inc.) and incubated for 1 min at room temperature. Grids were then washed with water (25 μl droplets) to remove excess graphene oxide. Pyrene-NTA solution (1.91 mM, 3 μl) was applied to the graphene oxide coated grid and incubated at room temperature for 5 min. Grids were further washed with water (50 μl droplets) before applying the ternary complex (250 μg/ml, 3 μl) followed by blotting for 6 sec and immediately plunge frozen into liquid ethane with the Cp3 Cryoplunger (Gatan, Inc.).

### Cryo-EM data collection and image processing

Data was collected using the Titan Krios (ThermoFisher) microscope equipped with a K2 Summit direct electron detector (Gatan) at Purdue Cryo-EM Facility (**Table 1** **and Fig. S5**). Sample grids were imaged at 300 kV, with an intended defocus range of 1.0 - 5.0 μm, a nominal magnification of 22,500X in super-resolution mode (0.668 Å per pixel), and at a dose rate of ~8 electrons per pixel per second (e.p.s.). Movies were collected with a total dose of ~45 electrons per Å^2^ (~1.12 electrons per frame per Å^2^) at 5 frames per seconds for 8 seconds. Of the total 1,731 movies collected, 1037 usable movies were aligned and dose weighted using MotionCor2 (21). CTF fitting was performed with Gctf (22). A total of 286,424 particles were picked using Gautomatch (Dr. Kai Zhang, MRC, UK). subsequent 2D and 3D classification, 3D refinement, post-processing, and local resolution estimation were performed in a beta release of Relion 2.0 (23). *De novo* initial model was constructed using EMAN2 (24).

**Table 1.** Cryo-EM data collection and refinement statistics

|  |  |  |
| --- | --- | --- |
| Number of grids used | 1 |  |
| Grid type | Lacey carbon and graphene oxide |  |
| Microscope/detector | Titan Krios/Gatan K2 |  |
| Voltage | 300 kV |  |
| Magnification | 22,500x |  |
| Recording mode | Super-resolution counting mode |  |
| Dose rate | 8 electrons per second per detector physical pixel |  |
| Defocus | 1.0 – 5.0 $\mu\text{m}$ | |
| Pixel size | 0.668 $\text{\AA}$/pixel | |
| Total dose | 45 $\text{e}/\text{\AA}^2$ | |
| Frame rate | 5 frame per second |  |
| Number of frames/movie | 40 |  |
| Total exposure time | 10 s |  |
| Number of micrographs | 1,731 |  |
| Number of micrographs used | 1,037 |  |
| Total particles picked | 286,424 |  |
| Number of particles used | 257,539 |  |
|  | **Holoenzyme** | **RPo** |
| PDB | 6C9Y | 6CA0 |
| EMD | EMD-7438 | EMD-7439 |
| Number of particles after 3D | 96,067 | 54004 |
| Particles used for final map | 96,067 | 54004 |
| Map resolution (FSC 0.143) | 4.2 $\text{\AA}$ | 5.75 $\text{\AA}$ |
| Local resolution (Resmap) | 4.1 $\text{\AA}$ -10 $\text{\AA}$ | 4.2 $\text{\AA}$ -10 $\text{\AA}$ |
| **Refinement (Phenix)** |  |  |
| Resolution ($\text{\AA}^2$) | 4.4 | 5.75 |
| Map CC | 0.768 | 0.838 |
| B factor ($\text{\AA}$) | 150.6 | 323.1 |
| **Validation** |  |  |
| All-atom clashscore | 6.67 | 9.99 |
| Rotamer outliers (%) | 5.32 | 4.20 |
| C-beta outliers | 5 | 0 |
| Overall score | 2.45 | 2.56 |
| **Ramachandran plot** |  |  |
| Favored (%) | 90.66 | 89.29 |
| Outliers (%) | 1.53 | 1.63 |
| **RMS deviation** |  |  |
| Bond length($\text{\AA}$) | 0.008 | 0.007 |
| Bond angle ($^\circ$) | 1.62 | 1.61 |

### Structure refinement

To refine the apo-form RNAP structure, *E. coli* RNAP holoenzyme crystal structure (PDB ID 4YG2) was manually fit into the cryo-EM density map using Chimera (25) and real-space refined using Phenix (26). In the real-space refinement, domains of RNAP were rigid-body refined, then subsequently refined with secondary structure, Ramachandran, rotamer and reference model restraints. To refine the structure of RPo, *E. coli* RNAP TIC crystal structure (PDB ID 4YLN) without RNA was manually fit into the cryo-EM density map using Chimera. Upstream DNA from the −35 element was manually built by using Coot (27). The structure was refined the same as in the apo-form RNAP.

### Electrophoretic Mobility Shift Assay

For testing the RNAP and DNA complex formation, 2,3,4 and 6 pmol RNAP was mixed with 4 pmol DNA (pFABC-3, **Fig. S2**) in binding buffer (10 mM Tris pH 8, 50 mM NaCl, 10 mM MgCl_2_, 5% glycerol, 0.01% Triton X100 and 1 mM DTT) and incubated at 37 °C for 10 min. RNAP-DNA complex was separated from free DNA by using 6% polyacrylamide-Tris-borate-EDTA (TBE) gel electrophoresis and DNA was visualized by ethidium bromide staining. The experiments were conducted twice.

### In vitro transcription Assay

*In vitro* transcription assays were performed in 10 μL volume containing 100 nM RNAP and 50 nM DNA (pFABC-3 or TI scaffold, **Fig. S2**) in transcription buffer (40 mM Tris-HCl (pH 8 at 25 °C), 30 mM KCl, 10 mM MgCl_2_, 15 μM acetylated BSA, 1 mM DTT) with nucleotide mixture (400 nM ATP, GTP and CTP plus 100 nM UTP and 1 μCi [γ-^32^P]UTP). The samples were incubated at 37 °C and the reactions were stopped by adding 10 μL of 2×stop buffer (90% formamide, 50 mM EDTA, xylene cyanol, and bromophenol blue). The reaction products were electrophoretically separated on a denaturing 24% polyacrylamide/7 M urea gel and visualized with a phosphorimager (Typhoon 9410; GE Healthcare). All experiments were conducted twice.

### 2-Aminopurine (2-AP) fluorescence assay

Synthetic DNA oligos (Integrated DNA Technologies) were obtained with 2-amino purine substituted at −7 or −1 positions of template strand DNA and annealed to the complimentary strand forming double stranded labelled DNA (pFABC_2AP-1 and pFABC_2AP-7, **Fig. S2**). 100 nM single stranded DNA, double stranded DNA and double stranded DNA mixed with 200 nM of purified wild-type RNAP or its σ^70^-R157E variant were incubated at 37 °C for 10 min, in a total of 100 μl reaction volume. The buffer composition of the mixture was kept as 10 mM Hepes pH 8, 50 mM NaCl, 5% glycerol, 1 mM MgCl_2_. Fluorescence signal from the samples were measured in a SpectraMax-M5 spectrophotometer (Molecular Devices) at excitation wavelength 320 nm and emission wavelength 380 nm.

## ACKNOWLEDGEMENTS

This work was supported by NIH grants GM087350 (K.S.M.) and GM093142 (D.Y.). We thank the Purdue Cryo-EM Facility (http://cryoem.bio.purdue.edu) for the use of the Titan Krios and CM200 microscopes.

## AUTHOR CONTRIBUTIONS

A.N, D.Y, W.J and K.S.M designed the research; F.V, K.L and A.N collected cryo-EM data; F.V, M.Z.Q, K.S.M and A.N analyzed the data; A.N and K.S.M wrote the paper. All authors discussed the results and commented on the manuscript.

